# The effect of cortical elasticity and active tension on cell adhesion mechanics

**DOI:** 10.1101/343038

**Authors:** B. Smeets, M. Cuvelier, J. Pešek, H. Ramon

**Affiliations:** KU Leuven – MeBioS, Kasteelpark Arenberg 30, 3001 Heverlee, Belgium

## Abstract

We consider a cell as a filled, elastic shell with an active surface tension and non-specific adhesion. We perform numerical simulations of this model in order to study the mechanics of cell-cell separation. By variation of parameters, we are able to recover well-known limits of JKR, DMT, adhesive vesicles with surface tension (BWdG) and thin elastic shells. We further locate biological cells on this parameter space by comparing to existing experiments on S180 cells. Using this model, we show that mechanical parameters can be obtained that are consistent with both Dual Pipette Aspiration (DPA) and Micropipette Aspiration (MA), a problem not successfully tackled so far. We estimate a cortex elastic modulus of *E_c_* ≈ 15 kPa, an effective cortex thickness of *t_c_* ≈ 0.3 µm and an active tension of *γ* ≈ 0.4 nN/µm. With these parameters, a JKR-like scaling of the separation force is recovered. Finally, the change of contact radius with applied force in a pull-off experiment was investigated. For small forces, a scaling similar to both BWdG and DMT is found.

Manuscript submitted to Biophysical Journal.

## Introduction

Cell adhesion is governed by a complex interplay between adhesive ligands and cortex that provides mechanical rigidity to the cell. Nonetheless, in controlled adhesion experiments with dual pipette aspiration (DPA) (1), it was found that classical JKR theory (2) for adhesion between solid elastic spheres is applicable. The pull-off force *F_s_* for separating a doublet of cells with effective radius 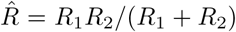 and adhesion energy w is close to the JKR formula, 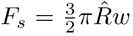, between the DMT formula 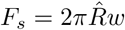, and the BWdG expression for adhesive vesicles with surface tension, 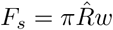 (1, 3).

Yet, there is an apparent mismatch between the elastic modulus estimated from such a pull-off experiment using JKR theory, and the Young’s modulus obtained from a simple single cell aspiration experiments (4). For example, on S180 murine sarcoma cells, a Young’s modulus^1^ *E_cell_* > 1 kPa is obtained from applying JKR theory to a DPA experiment (1), while single-cell aspiration tests (MA) on the same cell line yield a much lower modulus of *E_cell_* ≈ 100 Pa (4), even though both experiments are performed in similar conditions and at comparable timescales (seconds).

At first sight, an obvious reason for this is that the mechanical rigidity of a suspension cell is mostly concentrated in an elastic cortical shell rather than uniformly distributed throughout the cell. For thin elastic spherical shells, an expression for the pull-off force as a function of Young’s modulus *E_c_*, Poisson number *v_c_* and thickness *t_c_* has been derived in (5):
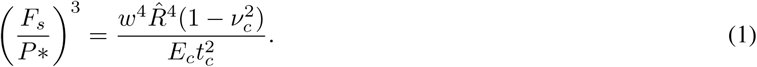

*P*\* is a dimensionless scaling factor that depends on the load conditions – for fixed load, *P\** ≈ 13.2 (5). A typical cell’s acto-myosin cortex has a thickness of roughly 200 nm and a Young’s modulus in the order of 10 kPa (6). Within this range of properties, and for a characteristic adhesion energy of *w* ≈ 1 nN/µm and a cell size of 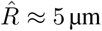, it can be verified using Eq. (1) that for a shell, the pull-off force 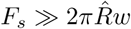, i.e. much greater than what was measured in (1). In other words, while the cell’s cytoskeletal structure resembles a thin shell, it behaves more like a solid elastic asperity during pull-off.

This discrepancy could be attributed to the highly non-linear and anisotropic behavior of the cortical acto-myosin and microtubule network (7). As such, its “effective” mechanical thickness would be significantly higher than the thickness measured using optical methods (6). This explanation is in line with observations of cortical rheology at long timescales, where a considerably elevated effective thickness is required to recover the rate of cell spreading using a simple Newtonian liquid model (8).

A second possible explanation lies in the active nature of the cortex: contractility induced by myosin II motors generates an active tension (*γ*), which counterbalances adhesion and thereby assists in the separation of two cells. For mature intercellular junctions, it has been shown that even the local regulation of contractility at the cell-cell interface rather than adhesion itself controls the extent of contact expansion. Then, the role of adhesion molecules is restricted to the mechanical anchoring of the cortex (9). While this local regulation of cortical tension is unlikely to affect adhesive behavior in controlled adhesion experiments at very short timescales (seconds), the total (uniform) cortical tension is likely to play a major role in a pull-off experiment. It should be noted here that this active tension is not the same as the surface tension often reported from mechanical tests (4, 6, 10), where a liquid model is used which assumes that no elastic stresses are present. This assumption can be valid at long timescales, when remodeling of the cortex effectively relaxes all elastic stresses. In absence of this relaxation, these two quantities would only coincide in the limit a of soft/thin cortex (see further).

Here, we propose a numerical model that tries to reconcile the aforementioned observations, describing adhesive contact between cells as a function of the elastic properties of the cell’s acto-myosin cortex and its active contractility. Cells are represented as spherical elastic shells that maintain internal volume and for which adhesive/repulsive contact is described using a Dugdale approach (11). Active contractility is explicitly introduced through a contractile tension, and acts similar to an additional surface tension (see Fig. 1). Using this model, we show how different scaling laws for a pull-off experiments can be recovered, by changing the stiffness *E_c_*, effective thickness *t_c_* and the active tension *γ* of the cortex. Next, in a case study on S180 suspension cells, we demonstrate how JKR-like behavior can be recovered during pull-off while remaining consistent to single-cell MA experiments. Doing so allows us to estimate the (instantaneous) mechanical properties of the S180 cell cortex.

**Figure 1:**
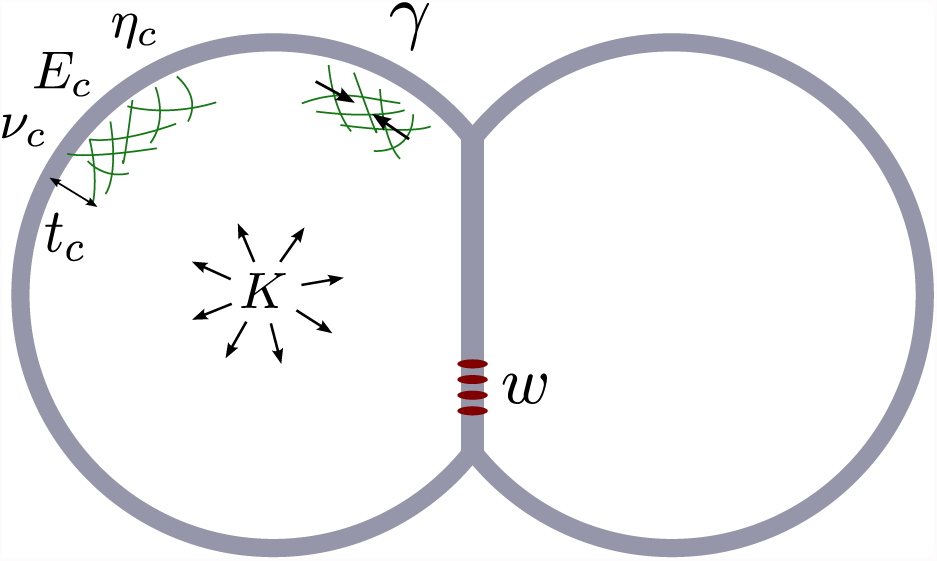
Schematic representation of a doublet of adhering cells: The cell’s cortex with thickness *t_c_* has passive elastic properties (Young’s modulus *E_c_*, Poisson ratio *v_c_* and viscosity *η_c_*) and an active contractile tension *γ*. Volume is maintained through a bulk modulus *K* and adhesion energy *w* drives the formation of cell-cell contacts.

## Methods

### Computational Model

We introduce a dynamical model that represents the cells as a triangulated spherical shell. An over-damped equations of motion 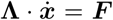 is solved to obtain the positions *x*, representing the node from a triangulated shell. Viscous (velocity-dependent) forces are represented in the resistance matrix **Λ**, whereas all other forces are assembled on the right-hand side in *F*. A shell model for linear visco-elasticity (Young’s modulus *E_c_*, Poisson number *v_c_*, thickness *t_c_* and viscosity η_c_) is implemented in a spring-damper network (see SI-1), where for simplicity we have assumed that *v_c_* = 1/3. To introduce active tension *γ* and conservation of volume with bulk modulus *K*, a local outward pressure is computed as:
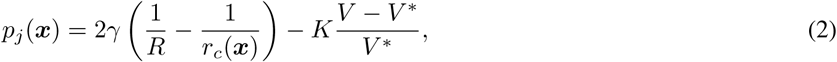

for a cell with radius *R*, volume *V*, and initial volume *V\**. *r_c_*(*x*) is the local mean radius of curvature on the cell surface at position *x*. We set *K* = 30 kPa, a value sufficiently high to prevent significant changes of cell volume during MA and DPA simulations. The cell’s surface is decorated with stickers, which are assumed to be fixed and uniformly distributed on the cell, and equal for both cells, leading to a work of interaction *w* = *w*_1_ + *w*_2_.

We aim to describe adhesive behavior in a wide range of cortical thickness. For larger *t_c_* and low *E_c_*, the normal (radial) elastic deformation of the cortex cannot be neglected anymore. Therefore, we use a modified Maugis-Dugdale contact model (11) that formulates a Hertzian repulsive pressure based on the contact stiffness, and an adhesive traction based on the adhesion energy w, and an effective range of interaction *h*_0_. For solid elastic spheres, the latter parameter captures the transition between the JKR (low *h*_0_) and the DMT limit (high *h*_0_) (12). For cells, the effective adhesive range is typically small and well in the JKR zone, and we set *h*_0_ = 50 nm (13). Since the Hertzian repulsive model is valid for a ‘solid’ elastic asperity, a requirement of this contact model is that the normal elastic compression of the cortex is small compared to its thickness. A discussion on the limitations of this model is provided in the ‘conclusion’ section, and a detailed description of the full computational methodology is presented in the SI-1-2.

### Simulation setup

Setups are created for numerical simulation of two mechanical tests: MA and DPA. For MA, we include an idealized pipette with a tip of toroidal shape of inner radius *R_p_* = 3.5 μm (4) – see Fig. 2(a) and SI. Within the pipette, an aspiration pressure *P_a_* is applied normal to the cell surface. For DPA, we first let two cells freely adhere until their contact area reaches a steady value – Fig. 2(b). Next, a pulling force is distributed – see SI – to the nodes of both cells (−*F_p_* and *F_p_*). We record the contact radius *R_c_* while the pulling force is gradually increased until the cells suddenly lose contact – Fig. 2(c). The force at which this occurs is registered as the separation force *F_s_*. Further details and numerical considerations of the simulation setup are provided in the SI.

**Figure 2:**
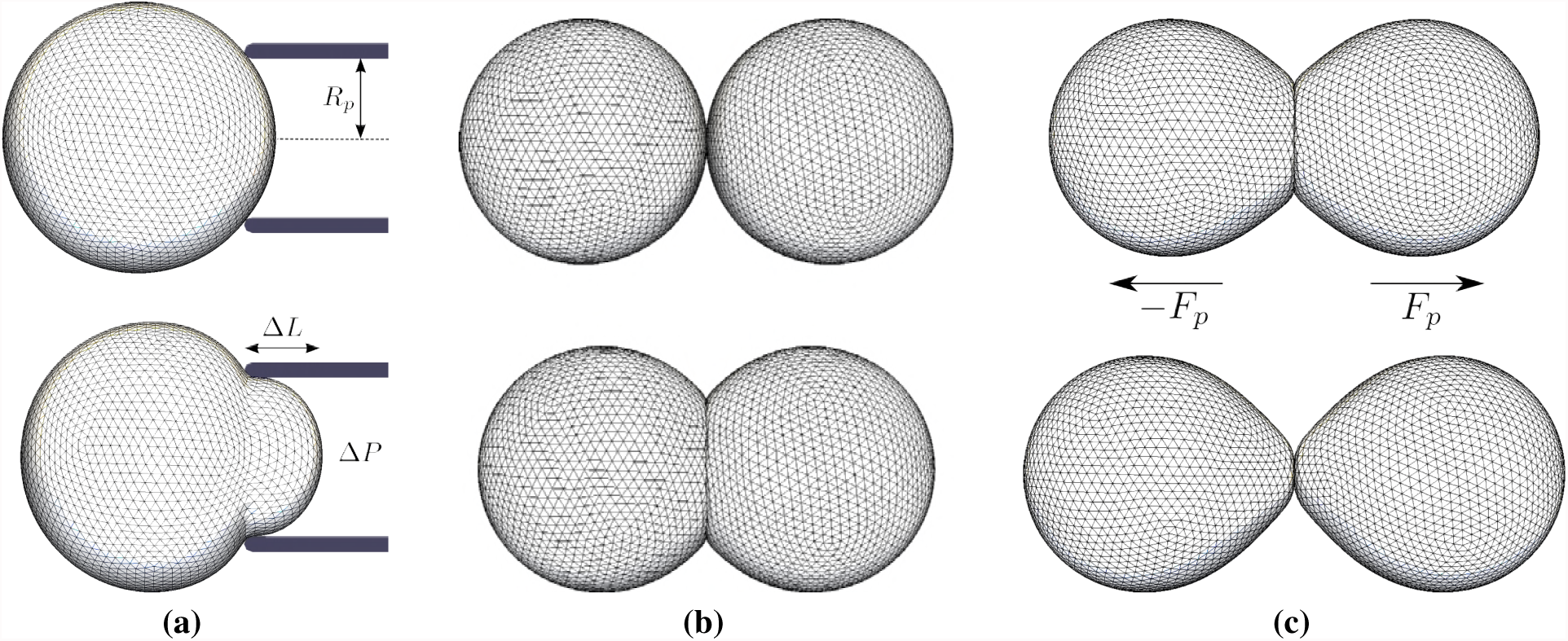
Visualization of simulation setups for MA and DPA experiments. **(a)**: Simulation of MA experiment. An underpressure Δ*P* is applied within a micropipette of radius *R_p_* (top). The pressure is gradually increased (bottom) until the aspirated length Δ*L* = *R_p_*. At this point, the pressure is registered as the critical pressure *P_c_*. **(b)**: Simulation of cell-cell adhesion. Two cells are brought in close proximity (top) and allowed to naturally adhere (bottom). **(c)**: Simulation of DPA experiment, starting from a doublet of adhering cells. A pulling force *F_p_* is applied on the cells (top) and gradually increased until rapid separation occurs (bottom). At this point, the pulling force is registered as the separation force *F_s_*.

## Results

### Pull-offforce in cell model

First, we show in general how the separation force depends on the mechanical properties of the cell’s cortex. For this, we define a dimensionless thickness as 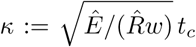. This chosen normalization is a ‘rigidity’ measure that ensures that a unique normalized separation force 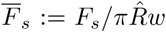 is found for the limits of BWdG 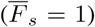, JKR 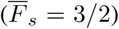, DMT 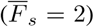 and shells – Eq. (1) – upon change of **κ**^2^. Here, we are mainly interested in the role of the effective cortical thickness *t_c_* and active tension *γ*.

Fig. 3 shows 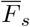 as a function of *κ* by varying *t_c_* for a simulated DPA experiment. Traversing from high *κ* (right) to low *κ* (left), four regions of distinct behavior can be recognized in these curves: (**I**) At high *κ*, or for contact radius *R_c_* ≪ *t_c_*, adhesion is dominated by localized elastic deformation normal to the contact plane. Here, solid-sphere Maugis-Dugdale adhesion is recovered, and 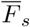 will range from the JKR to the DMT limit. (**II**) for lower *κ*, or 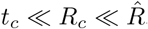, the contribution of bending resistance is dominant (bending rigidity 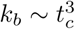). This resistance to curvature change is similar to surface tension: the BWdG limit 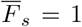 is approached. (**III**) as *κ* decreases, a sharp increase in 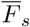 is observed, similar to shell theory. The adhesion energy is balanced by in-plane elastic energy distributed over the complete cortex. (**IV**) At very low *κ*, the complete cortex is under high strain, and shell theory breaks down. A maximal contact radius *R_c_* in the order of 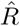 is reached and volume conservation (bulk modulus *K*) limits further deformation. A plateau is observed at large values of 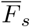. It should be noted that the proposed computational description becomes invalid at both extremes of *κ*. At very large *κ*, the thin-shell assumption of cortex elasticity breaks down. At low *κ*, indentations will become large compared to thickness, and the assumptions of our adhesion model break down. While this can be safely mitigated by replacing the normal contact stiffness with a sufficiently stiff constraint, the system becomes prone to buckling instabilities in the absence of active tension. Unsurprisingly, the role of active tension γ is mainly significant at small values of *κ*, where it reduces 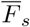 towards the JKR-DMT zone. Given what we know about typical mammalian cells (see introduction), we expect *κ* to be small, even if an ‘effective’ thickness would be much greater than the optical thickness. In this case, a significant cortical tension is required for a pull-off force to be in the JKR-DMT zone, 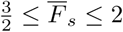.

**Figure 3:**
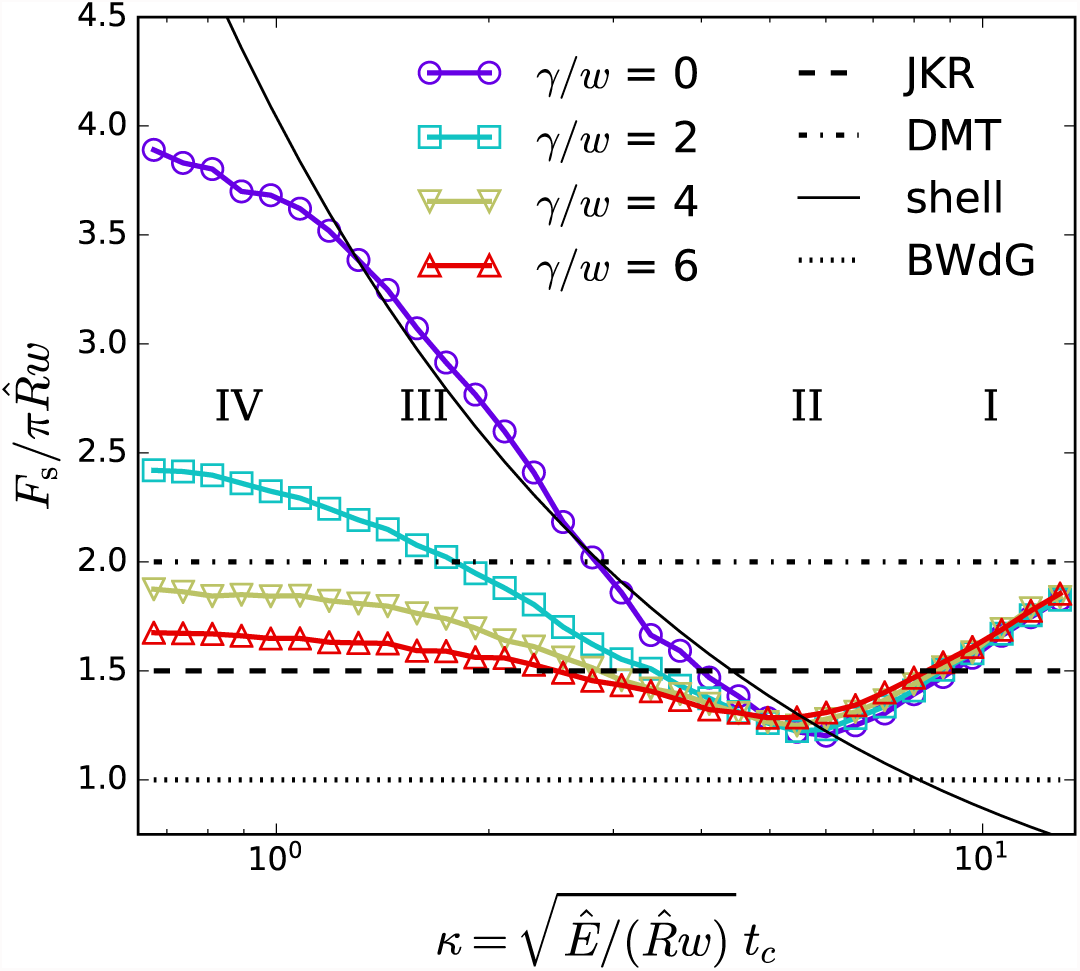
Normalized pull-off force 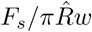 as a function of dimensionless thickness *κ* for varying active tension *γ*. The simulations were obtained by varying thickness *t_c_*, while 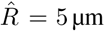, *E* = 25 kPa and *w* = 0.25 nN/µm. For reference, the pull-off force from JKR, DMT, BWdG and shell theory are shown for these parameters.

### Case study on S180 cells

We try to locate cells in this general framework by considering S180 cells, a mechanically very well investigated cell line. In Chu et al. (2005) (1), pull-off forces were measured using DPA in controlled adhesion experiments. We have replotted these results in Fig. 4(a). A scaling of 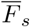 in the region of the JKR and DMT limits can be observed, with an average 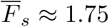 in the sampled region of *w*. In our model, such a scaling could be obtained for many possible combinations of *E_c_*, *t_c_* and *γ*. To restrict the parameter space to realistic cell properties we first compare to separate MA experiments on the same cell line. In Engl et al. (2014) (4), a (liquid model) mean cortical tension of 0.9 nN/µm was found^3^ from MA on S180 cells, which corresponds to a critical pressure *P_c_* ≈ 250 Pa. We sampled combinations of *E_c_*, *t_c_*, and *γ* in a full factorial 15 × 15 × 15 grid and performed MA simulations to compute the critical pressure (see SI-2). From this, an iso-surface was extracted that represents all parameter combinations yielding a critical pressure of 250 Pa – Fig. 4(b). Subsequently, we re-sampled points in a regular distribution on this iso-surface. Each of these points represents a combination of *E_c_*, *t_c_*, and *γ* that resembles the mechanical behavior of an (average) S180 cell in a MA experiment.

**Figure 4:**
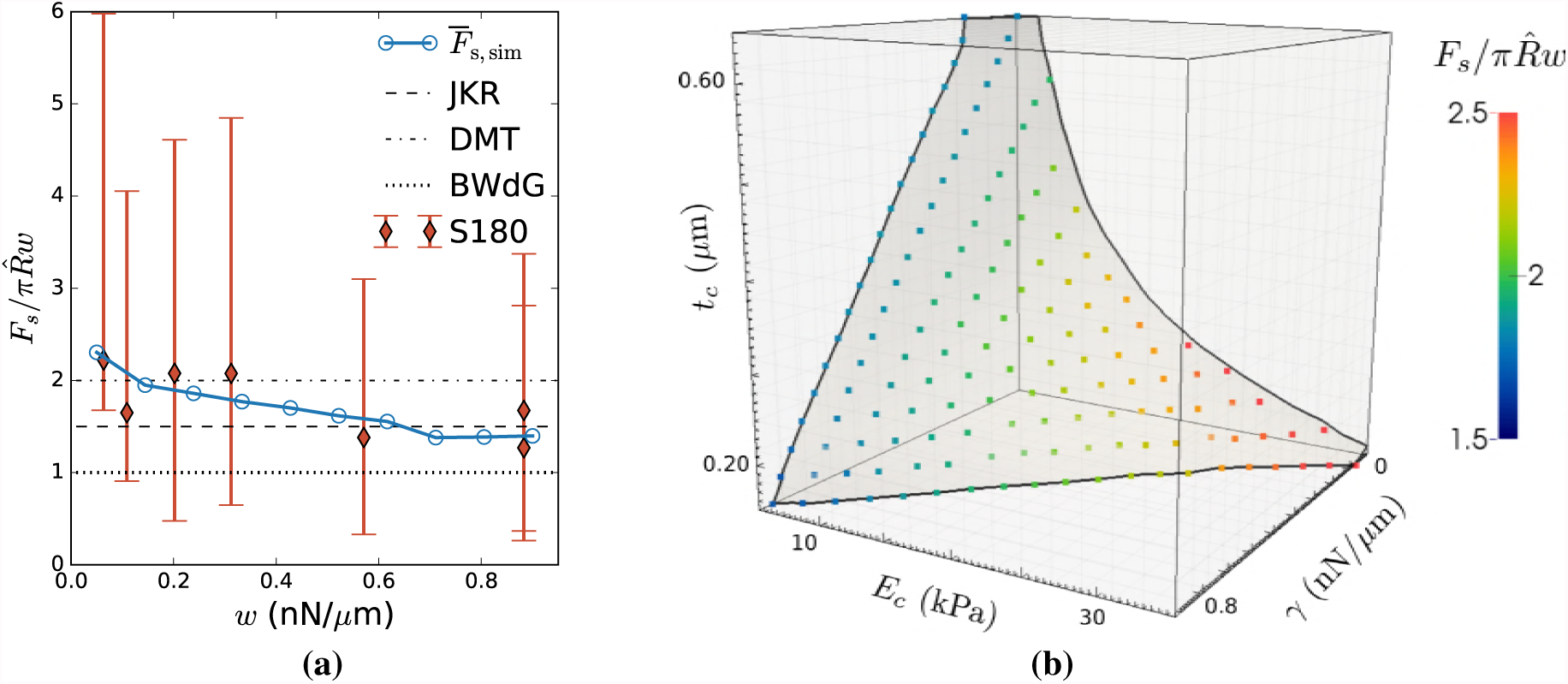
**(a)** Normalized DPA pull-off force 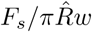 as a function of dextran depletion-induced adhesion *w* between S180 cells, replotted from Fig. 2 in Chu et al. (2005) (1) (red diamonds), together with simulated DPA experiment at *E_c_* = 15 kPa, *t_c_* = 0.3 µm, and *γ* = 0.4 nN/µm, as consistent with MA data (blue circles). Guide-lines with BWdG, JKR and DMT limits are provided as indication. We have assumed R = 6 µm **(b)** Parameter space of *E_c_*, *t_c_* and *γ*, with an iso-surface obtained from simulations of MA on S180 cells, which delimits all parameter combinations for which the experimentally observed critical pressure *P_c_* = 250 Pa (4) is attained. Colored dots represent new samples in this surface for which simulations of DPA were performed. The color scale indicates the value of 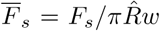 obtained for an adhesion energy of *w* = 0.5 nN/µm. The overlaid rectangular lattice depicts the grid in which MA simulations were performed.

Finally, we performed simulations of DPA on these new samples at an intermediate *w* = 0.5 nN/µm, and registered 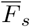. The result of this can be seen in Fig. 4(b). The lower values of 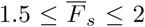 observed in (1) occur only at lower *E_c_*, when additional resistance to deformation is offered by either bending rigidity (at higher *t_c_*) or active tension *γ*. This confirms our hypothesis that either the presence of active tension, or a larger ‘effective thickness’ is required to explain the adhesive behavior of cells. Moreover, it can be observed that a manyfold increase of *t_c_* is required to have the same effect as a moderate active tension. Under the assumption that the apparent increase in *t_c_* is moderate, *t_c_* = 0.3 µm, we estimate for S180 cells that *γ* ≈ 0.4 nN/µm and *E_c_* ≈ 15 kPa. The full parameter set of estimated properties is listed in Table 1. It should be stressed that the goal of this work is not to determine the mechanical properties of S 180 cells, but rather to demonstrate a quantitative relationship obtained between *E_c_*, *t_c_* and *γ*, and provide an estimate for the range of possible parameters.

**Table 1:** Table of numerically estimated mechanical properties of S 180 cells that is consistent with MA and DPA experiments.

| Parameter | Symbol | Value | Unit |
| --- | --- | --- | --- |
| Young’s modulus cortex | $E_c$ | 15 | kPa |
| Poisson’s ratio cortex | $\nu_c$ | 1/3 | - |
| Thickness cortex | $t_c$ | 0.3 | $\mu\text{m}$ |
| Active cortical tension | $\gamma$ | 0.4 | $\text{nN}/\mu\text{m}$ |
| Bulk modulus cell | $K$ | 30 | kPa |
| Cell radius | $R$ | 6 | $\mu\text{m}$ |
| Effective adhesive range | $h_0$ | 50 | nm |

We performed simulations of DPA with the parameters from Table 1 for varying adhesion energy and overlay the resulting 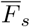 with experimental values from (1) – see Fig. 4(a). A close correspondence is found between simulation and experiment, and in both cases, a small decrease of 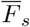 with *w* is observed, with 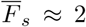 for small *w* and 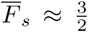 for larger *w*. This trend is similar to the transition observed for solid elastic spheres: *w* affects the Tabor parameter 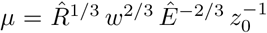 (with effective range of interaction *h*_0_ ≈ 0.97 *z*_0_) (12) that describes the transition between JKR-like and DMT-like adhesion.

While we were not able to formulate a similar universal transition parameter for our more complex modeled system, the underlying mechanisms can be similar: At low *w*, *h*_0_ is large compared to the contact radius, and the region of adhesive traction is affected little by elastic deformation (DMT-assumption). At large *w*, the elastic deformation is much greater than *h*_0_, and fully determines the adhesive region (JKR-assumption).

### Contact radius

For the parameters in Table 1 and *w* = 0.6 nN/µm (hence, *κ* ≈ 0.87), we investigate in detail the change of contact radius *R_c_* with increase of the applied pulling force *F* in a DPA experiment. For solid elastic spheres, the contact radius depends on the elastic modulus 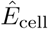. For DMT and JKR, the cube of the contact radius are given by:
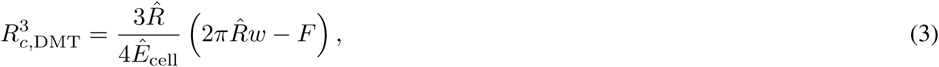

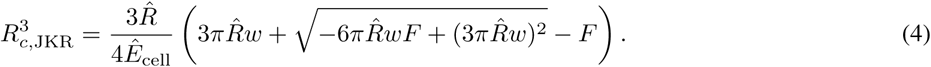

Further, we will define the relative contact radius *ψ* := *R_c_*/*R*. For the BWdG model for adhesive vesicles with surface tension, the relationship between applied pull force and contact radius is non-trivial and is expressed in function of the deformed apex radius *R_a_* (3). With *ψ*′ := *R_c_*/*R_a_*:

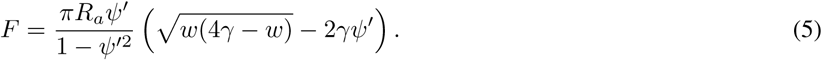

Fig. 5 shows *ψ* and *ψ*′ as a function of F for a simulated cell, compared to the theoretical predictions of DMT, JKR and BWdG. For DMT and JKR, we expect the change of apical radius to be small so approximate *R_a_* ≈ *R* hence *ψ*′ ≈ *ψ*). From this comparison, we list the following observations:

I. Pull-off force is close to the JKR limit. The maximal contact radius at F = 0 corresponds to an apparent elastic modulus – obtained from Eq. (4) – 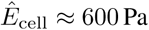. However, the change of contact radius with force does not follow JKR theory. The dependency of the effective 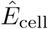 on *γ* is further shown in SI Fig. S4.
II. Rupture occurs at much higher tensile loading than for ideal adhesive vesicles with surface tension (BWdG) due to the presence of bending resistance that ensures the maintenance of low contact angles. The maximal contact radius at *F* = 0 corresponds to an adhesive vesicle with a surface tension of ≈ 0.8 nN/µm, which is in close agreement with the value of 0.9 nN/µm obtained from the analysis of MA experiment, assuming that the cell is a liquid droplet with surface tension (4). This correspondence of cell contact radius to the BWdG model in (self- or externally) compressed conditions but not during tensile loading and pull-out has been observed experimentally on HeLa cells in (14).
III. At low *F* (large contacts) the change of *ψ* with *F* is DMT-like, i.e. 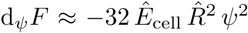, with an apparent 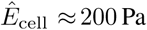 when using Eq. (3). This indicates that the force-indentation *F*(*δ*) response of adherent cells with surface tension is remarkably Hertzian. This observation was confirmed in a simulated compression test on a spread out cell: Around *F* = 0, the contact force follows *F* ~ *δ*^3^/^2^ (see SI Fig. S5). This contrasts with the force-deformation response of a liquid-filled shell with no adhesion or active tension, which showcases cubic behavior *F* ~ *δ*^3^ (15).

**Figure 5:**
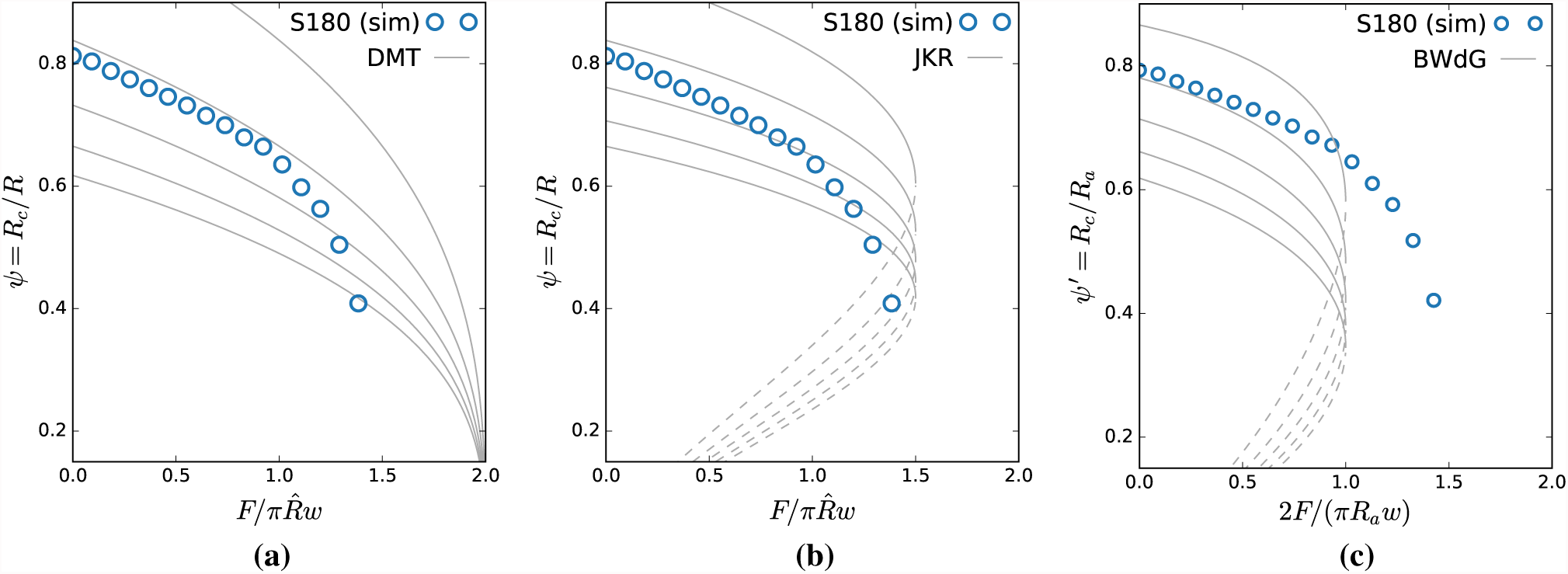
**(a)** Normalized contact radii *ψ* and *ψ*′ as a function of normalized applied force 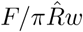 of a simulated S180 DPA pull-off experiment (parameters in Table 1) and for *w* = 0.6nN/µm, compared to (**a**) DMT, (**b**) JKR and (**c**) BWdG limits. For solid elastic spheres (JKR and DMT), we assume that *R_a_* ≈ *R*. Dashed lines indicate where contact is not stable, and rupture of the cell doublet will occur. Different guide lines are shown (from top to bottom at *F* = 0) for DMT: 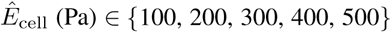, for JKR: 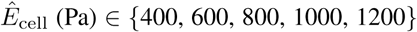 and for BWdG: *γ* (nN/µm) ∊ {0.6, 0.8, 1.0, 1.2, 1.4}.

## Conclusion

In this work, we have quantified the adhesion behavior of filled elastic shells with active tension, which were used as a model for biological cells. Numerical simulations with this model were carried out in order to investigate the role of cortical stiffness, thickness and active tension. These simulations showed that a combination of these properties can simultaneously explain the mechanics of cell deformation during aspiration and of cell-cell separation during a pull-off experiment. We estimate that cells exist in small to moderate ranges of a dimensionless thickness *κ* In these conditions, the active tension plays a crucial role, and is required to explain the observed scaling of separation force.

By comparing to existing experiments on S180 cells, we give tentative estimates of mechanical properties of their actin cortex. These estimates are in good agreement with various other characterizations in literature (1, 4, 6). We show that JKR-like pull-off forces can exist for a wide range of adhesion energies. Secondly, the scaling of contact radius with force at low loading force (or, conversely, large adhesive deformation) follows BWdG predictions, but is also consistent with DMT theory, implying that force-deformation behavior is Hertz-like at sufficient deformation. This suggests that in an indentation experiment, for example Atomic Force Microscopy (AFM), this model would be almost indistinguishable from a solid elastic material. Since Hertz theory is ubiquitously used in AFM experiments to parameterize cells with an apparent Young’s modulus, this begs the question of where its application is appropriate and where not. It can be argued that for strongly spread out cells, where dense cytoskeletal material spans the full height of the cell, this parameterization is apt, but that for more rounded cells (e.g. monocytes), an approach more similar to (6) is advisable.

### Limitations

We have presented a minimalistic mechanical model that disregards most of the complexities that accompany cell-cell adhesion in real biological settings. Some of these complications can well be expected to affect the results presented here in a non-trivial manner, and they will be briefly discussed here.

Firstly, we considered the cell’s cytosol as liquid-like (i.e. bearing hydrostatic stresses, and this through an effective bulk modulus *K*). The physical properties of the cell’s internal structures are complex, and models that capture its mechanical behavior are often dependent on the timescale of interest. At fast timescales (≪ 1 s), indented cells show a viscoelastic creep response, which might be attributed to Maxwell fluid behavior of the cell’s internal structures (16). Hence, the assumption in our analyses was that experiments were at least slow enough to relax any deviatoric stresses.

Secondly, our shell model consists of linearly elastic material, while the cell cortex has been shown to exhibit non-linear behavior at large deformations, including both strain stiffening and strain softening (7). Typical strains in our simulated experiments are very low (< 5%), but can locally reach up to 20% e.g. near rupture at the contact site in a pull-off experiment. Here, non-linearities in stretch response could have non-negligible effect on the separation force.

Finally, we model adhesion based on the assumption of fixed and non-specific stickers (or, with a mobility timescale that is much slower than the timescale of bond rupture). This assumption is valid for the experiment we compared against, where depletion-induced adhesion was studied, but (17). In naturally adhering cells, adhesive ligands have been shown to diffuse in the plasma membrane and cluster at the site of cell-cell junctions (18). All these phenomena are expected to affect adhesion and debonding mechanics, both dynamically and at steady-state. While not the focus of this study, these properties need to be taken into account to model cell-cell adhesion in realistic biological settings.

## Author Contributions

B.S. designed research; B.S., J.P. performed research; and B.S., J.P., M.C. and H.R. wrote the paper.

## Acknowledgements

We thank S. Vanmaercke for technical support in designing the computational algorithms for the numerical simulations. B.S. acknowledges support from the Research Foundation - Flanders (FWO), Grant Nr. 12Z6118N.

## SUPPLEMENTARY MATERIAL

An online supplement to this article can be found by visiting BJ Online at http://www.biophysj.org.

## Footnotes

1 Only the composed modulus 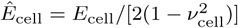 can be estimated this way.

2 However, we do not expect identical 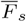. from simulations of our model for all parameter combinations that lead to the same *κ*.

3 This value of 0.9 nN/µm already gives an upper limit for estimates of the active tension *γ* which should be approached in the limit of a soft and thin elastic shell.

